# The transcription factor NF-Y promotes myeloid cell survival and protects from inflammatory vascular disease

**DOI:** 10.64898/2026.01.29.702697

**Authors:** Carlos Silvestre-Roig, José M González-Granado, Pilar Gonzalo, Lina-Marie Vöcking, Raphael Chevre, Vanesa Esteban, Vicente Andrés

**Affiliations:** Institute of Experimental Pathology, Center for Molecular Biology of Inflammation (ZMBE), University of Münster, Germany; Centro de Investigación Biomédica en Red de Enfermedades Cardiovasculares (CIBERCV), Madrid, Spain; LamImSys Laboratory, Instituto de Investigación Sanitaria Hospital 12 de Octubre (imas12), Madrid, Spain; Department of Immunology, Ophthalmology and ENT, School of Medicine, Universidad Complutense de Madrid (UCM), Madrid, Spain; Centro Nacional de Investigaciones Cardiovasculares (CNIC), Madrid, Spain; Department of Allergy and Immunology, IIS-Fundación Jiménez Díaz, UAM, Madrid, Spain; Faculty of Biomedical and Health Sciences, Universidad Alfonso X El Sabio, Madrid, Spain

**Keywords:** NF-Y, atherosclerosis, endoluminal femoral artery injury, macrophage, neutrophil

## Abstract

**Background:** Myeloid cells orchestrate vascular inflammation through transcriptional programs that regulate their maturation, effector function, and survival. While lineage-determining transcription factors establish myeloid identity, understanding of the transcriptional regulation of myeloid behavior in chronic inflammatory contexts remains limited. Nuclear factor-Y (NF-Y) is a trimeric CCAAT-binding transcription factor that regulates cell proliferation and differentiation. Here, we investigate the role of NF-Y in myeloid function and survival during chronic vascular inflammation.

**Methods:** Integrated single-cell transcriptomics of BM, blood, and atherosclerotic lesions were combined with myeloid-specific NF-YA inactivation to define NF-Y–dependent transcriptional states. Functional consequences were assessed in mice with myeloid-specific *Nfya* deletion on a hypercholesterolemic *Apoe*^*–/–*^ background using models of diet-induced advanced atherosclerosis and endoluminal femoral injury. Myeloid cell recruitment, survival, apoptosis, and proliferation were further examined in models of thioglycolate-induced peritonitis.

**Results:** NF-Y subunit transcripts were detected across myeloid compartments, with *Nfya* enriched in proliferative macrophages and immature neutrophils. In mouse atherosclerotic lesions, low *Nfya* expression was associated with lipid-handling and phagocytic macrophage signatures and a pro-inflammatory neutrophil phenotype. Myeloid *Nfya* deficiency was further associated with reduced circulating neutrophil counts, increased macrophage and neutrophil apoptosis during acute inflammation, expanded necrotic cores, larger unstable atherosclerotic lesions, and aggravated atherosclerosis and injury-induced neointimal thickening.

**Conclusion:** Our data identify NF-Y as a transcriptional safeguard of myeloid cell survival during inflammatory stress, thereby shaping disease progression and outcomes in vascular disease.

## Introduction

Myeloid cells are key determinants of inflammation during vascular disease progression and its associated complications, including myocardial infarction and stroke^1^. In atherosclerosis, monocytes and macrophages drive lesion growth through lipid uptake and amplification of inflammatory cascade, while neutrophils contribute to early recruitment events and promote plaque instability through tissue injury and sustained inflammation^2^. These pro-inflammatory activities arise from the activation of transcriptional programs both at sites of myeloid cell production and locally within inflamed vascular tissue. Transcriptomic and epigenomic studies have identified the transcription factors that—beyond lineage specification—control the activity of macrophages and neutrophils in these contexts.

Macrophage functions are regulated by networks of transcription factors that integrate external signals to shape gene expression programs underlying activation, polarization, and homeostatic maintenance^3^. Several lineage-determining and stimulus-responsive transcription factors orchestrate macrophage identity and functional phenotypes in response to cytokines and environmental cues, including Signal Transducer and Activator of Transcription (STAT), Nuclear Factor-κB (NF-κB), Interferon Regulatory Factors (IRFs), Peroxisome Proliferator-Activated Receptors (PPAR), and Kruppel-like Factors (KLF). Similarly, neutrophil transcriptional programs are controlled by distinct transcription factors that regulate differentiation and maturation in the bone marrow (BM) (e.g. Runx1, Klf6), mobilization to the circulation, recruitment to tissues, and effector functions (e.g. Junb, Relb)^4^. Together, these findings highlight a central role for transcriptional regulators that maintain myeloid proliferative capacity, stress tolerance, survival, and effector function, extending beyond the establishment of lineage identity.

Nuclear factor-Y (NF-Y) is a trimeric transcription factor composed of NF-YA, NF-YB, and NF-YC subunits that binds to CCAAT motifs commonly found in promoters of genes involved in core cellular processes, including cell-cycle progression and biosynthetic pathways^5^. In hematopoiesis, NF-Y has been linked to stem and progenitor cell function, maintaining cell survival in proliferative—but not quiescent—states.^6, 7^ Consistent with this role, overexpression of a dominant-negative form of NF-YA reduces myeloid and granulocyte progenitor populations, impairing cell expansion without blocking differentiation.^8^ Mechanistically, NF-Y regulates G2/M phase exit through interactions with the master regulator CCAAT/enhancer-binding protein ε (CEBPε), acting via inhibition of E2F and NF-Y activity to initiate neutrophil differentiation^9^.

Beyond its role in proliferation control, NF-YA expression increases during monocyte-to-macrophage differentiation^10^ and modulates inflammatory responses upon myeloid cell activation.^11, 12^ These observations position NF-Y not only as a regulator of cell-cycle progression and survival, but also as a potential integrator of proliferative inflammatory programs in myeloid cells. Here, we combine integrated single-cell transcriptomic analysis across BM, blood, and atherosclerotic lesions with genetic inactivation of NF-YA in myeloid cells to interrogate the role of NF-Y in myeloid cell survival and inflammatory responses. Using mouse models of chronic vascular disease and acute inflammation, we demonstrate that loss of NF-Y disrupts myeloid cell survival, enhances inflammatory pathology, and accelerates vascular disease progression.

## Methods

### Data availability

All data supporting the findings of this study are available within the article, the scRNA-seq data is available under GSE316606, and the code used for scRNA-seq analysis is deposited in GitHub (https://github.com/csilvestreroig/Silvestreroig_lab.github.io/tree/main/projects/NF-Y_project).

### Mouse models and animal procedures

All animal procedures were conducted in accordance with institutional guidelines and regulations and were approved by the local Ethics Committee. Experiments were conducted on female and male mice.

### Mouse strains and genotypes

*Nfya*^*fl/fl*^ mice (aka *Cbfa*^*flox/flox*^) on a C57BL6/J background were as previously described.^13^ These mice were crossed with *LysM*^*CRE/+*^ mice to generate myeloid-specific *Nfya* deletion and were further backcrossed onto an apolipoprotein E (*Apoe*^−*/*−^) background for atherosclerosis studies. Unless otherwise indicated, control mice with intact *Nfya* were *LysM*^*+/+*^ *Nfya*^*fl/fl*^ littermates on the same genetic background. Mice were maintained on standard chow diet (2.8% fat; Panlab, Barcelona, Spain) unless specified.

### Bone marrow transplantation and high-fat diet-induced atherosclerosis

For BM transplantation, donor and recipient mice were maintained on an *Apoe*^−*/*−^ background to ensure equivalent systemic lipid profiles across experimental groups. Recipient female *Apoe*^−*/*−^ mice were lethally irradiated with two doses of 6.5-Gy administered 3 hours apart (JL Shephed & Associates 1-68A irradiator, 1000 Curie ^137^Cs source). After 24 hours, mice received 2 × 10^6^ BM cells by intravenous injection. Antibiotics were administered for 7 days before and after irradiation. After a 4-week hematopoietic reconstitution period, atherosclerosis was induced by placing mice on a high-fat diet (HFD: 10.8% total fat, 0.75% cholesterol; S8492-E010, Ssniff, Germany) for 12 weeks.

### Femoral artery wire injury

Endoluminal injury of the common femoral artery was performed in 2-month-old male and female *Apoe*^−*/*−^ *LysM*^*CRE/+*^ *Nfya*^*fl/fl*^ mice and *Apoe*^−*/*−^ *Nfya*^*fl/fl*^ controls maintained on chow diet. A 0.25-mm diameter guidewire (Advanced Cardiovascular Systems) was introduced into the femoral artery, advanced to the aortic bifurcation, and withdrawn as described.^14^ Mice were sacrificed 7 days after injury, and tissues were cleaned and fixed in 4% paraformaldehyde for subsequent analysis. Percent stenosis was calculated as

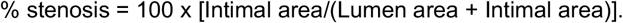

### Thioglycolate-induced peritonitis

Acute inflammatory studies were performed with male and female *LysM*^*CRE/+*^ *Nfya*^*fl/fl*^ mice and *Nfya*^*fl/fl*^ controls on a C57BL/6J background (with intact *Apoe*). Acute peritonitis was induced by intraperitoneal injection of 3% thioglycolate. Mice were sacrificed 16 hours after injection, and peritoneal exudates were collected by intraperitoneal lavage with 10 mL of cold PBS following gentle abdominal massage.

For competitive repopulation experiments, lethally irradiated recipient male and female mice were reconstituted with a 1:1 mixture of RFP^+^ BM cells isolated from dsRED reporter mice and RFP^−^ cells isolated from *LysM*^*CRE/+*^ *Nfya*^*fl/fl*^ mice. After hematopoietic reconstitution, thioglycolate-induced peritonitis was performed as described above.

### Single-cell transcriptomic studies

For single-cell RNA sequencing, *Apoe*^−*/*−^ female mice were fed a HFD for 16 weeks. Mice were then euthanized, and blood, BM, and brachiocephalic arteries were collected for cell isolation and downstream processing as described below.

Single cells suspensions from blood, BM and atherosclerotic tissue were quantified by flow cytometry, normalized across conditions, and labeled with hashtag antibodies specific to each tissue of origin. After extensive washing, cells were pooled and fluorescence-activated cell sorted (FACS) as DAPI^-^ CD115^+^ Ly6C^+^ from blood and BM, and as DAPI^-^ CD45^+^ from aortic tissue. Sorted cells were loaded onto the 10x Genomics platform and processed using the Chromium Next GEM Single-Cell 3′ Reagent Kit with Feature Barcoding technology. Libraries were sequenced by Novogene. Raw sequencing reads were processed with Cell Ranger v6.0.2 and aligned to the *Mus Musculus* reference genome (GRCh38 mm10).

Single-cell RNA-seq datasets were obtained from publicly available 10x Genomics–formatted count matrices and from the in-house dataset described above. Public datasets comprised multiple mouse atherosclerosis- and aorta-related studies: Cochain et al.^15^ (GSE97310), Lin et al.^16^ (GSE123587), Williams et al.^17^ (GSE154817), Cyr et al.^18^ (GSE252243), Kim et al.^19^ (GSE116240), and Amoros et al.^20^ (E-MTAB-15242), as well as an additional single-sample dataset (GSM4981311). For each dataset, sparse UMI count matrices were read using Matrix::readMM, and barcode and feature tables were imported using data.table::fread. A SingleCellExperiment (SCE) object was generated for each dataset, with raw counts stored in the counts assay. Cell barcodes were prefixed with dataset-specific identifiers to enable tracking of sample origin.

To integrate datasets across studies, gene identifiers and metadata were harmonized. Within each dataset, duplicated gene symbols were collapsed by summing counts to yield a single row per unique gene symbol. The intersection of gene symbols common to all datasets was then determined, and each dataset was subset to this shared gene set. Row names were replaced with canonical gene symbols, and non-count assays and reduced dimension representations were removed prior to merging.

Genes with zero total counts across the merged dataset were removed. Libraries were size-normalized and log-transformed using scuttle::logNormCounts. Highly variable genes (HVGs) were identified with scran::modelGeneVar, and HVGs were selected using scran::getTopHVGs with proportion-based selection. Principal component analysis (PCA) was performed on HVGs using scater::runPCA, with results stored as reduced-dimension embeddings. To mitigate between-study batch effects, Harmony^21^ was applied to the PCA space using harmony::HarmonyMatrix, with *Condition* specified as the batch variable. Uniform Manifold Approximation and Projection (UMAP) embeddings were computed from Harmony-corrected coordinates using scater::runUMAP, based on the top 10 dimensions. A shared nearest-neighbor (SNN) graph was constructed on the corrected PCA space using scran::buildSNNGraph (k = 30), and community detection was performed by Louvain clustering (igraph::cluster_louvain). Cluster labels were stored as colLabels in the SCE object.

Cell types were annotated by reference-based label transfer using SingleR (SingleR::SingleR) with the ImmGen reference dataset (celldex::ImmGenData). Reference marker genes were summarized per cluster, and clusters were assigned to broad cell classes based on SingleR predictions combined with curated inspection.

Per-cell quality-control metrics were computed using scater::perCellQCMetrics and scater::addPerCellQC. Mitochondrial genes were identified using a prefix match (^mt-). Low-quality cells were flagged as outliers based on library size and number of detected genes (lower outliers identified on log-transformed values using a median absolute deviation–based approach) and by applying a mitochondrial transcript proportion threshold (>5%). Cells meeting any of these criteria were excluded. Feature-level filtering was then applied: genes not expressed in any cell were removed, low-abundance genes were filtered based on average expression, and genes expressed in ≤5 cells were excluded.

After subsetting of neutrophils or macrophages, marker genes defining clusters were identified. Genes were first filtered to those expressed in a minimum fraction of cells within each group, defined as the proportion of cells per cluster with expression above a specified threshold. Differential expression analysis was performed on raw counts using edgeR (edgeR::DGEList, calcNormFactors). A no-intercept design matrix (∼0 + label) was used, dispersions were estimated (estimateDisp), and quasi-likelihood negative binomial models were fitted (glmQLFit, robust = TRUE). Pairwise contrasts between clusters were tested with glmTreat, applying a minimum log_2_ fold-change threshold of log_2_(1.5). Differential expression results were ranked and used to construct cluster-specific gene signatures (top-N genes per contrast) using the CreateGeneSignatures package (https://github.com/ATpoint/CreateGeneSignatures).

*Nfya*^*high*^ and *Nfya*^*low*^ cells were defined based on log-normalized *Nfya* expression. Multiple thresholds were evaluated (median, 75th, 90th, and 95th percentiles); primary analyses used the 95th percentile cutoff (top 5% of cells) to define *Nfya*^*high*^ cells (*Nfya_q95*), with all remaining cells classified as *Nfya*^*low*^. Groups assignments were visualized on UMAP embeddings and validated using expression histograms. For *Nfya*^*high*^ versus *Nfya*^*low*^ comparisons, genes were filtered to those expressed in at least a minimum fraction of cells in either group. Differential expression was assessed using edgeR quasi-likelihood models with a design matrix defined by *Nfya* expression status (∼0 + *Nfya_q95*), and contrasts were tested using glmTreat with a log_2_ fold-change threshold of log_2_(1.5). Volcano plots were generated from the resulting differential expression tables (excluding *Nfya*), highlighting the most strongly differentially expressed genes.

All analyses were performed in R using Bioconductor and CRAN packages, including but not limited to SingleCellExperiment, scater, scran, edgeR, Harmony, SingleR/celldex, escape, ComplexHeatmap, clusterProfiler, and ReactomePA.

### Signature scoring and functional enrichment analysis

Gene set scores (**Table_01**) were computed using ESCAPE (escape::runEscape) with the UCell method^22^ on trimmed gene sets (maximum of 1500 genes per set). The resulting score matrix was added to the colData slot for downstream analyses. For each gene signature, differences between *Nfya*^*high*^ and *Nfya*^*low*^ were assessed using Welch’s two-sample *t*-test (rstatix::t_test), excluding signatures with insufficient variability or missing values. Multiple testing correction across signatures was performed using the Benjamini–Hochberg false discovery (FDR) method. Signature score distributions were visualized using violin and box plots, and summary visualizations (e.g., lollipop plots) were generated from mean score fold changes (*Nfya*^*high*^ / *Nfya*^*low*^) across categories.

Functional enrichment of differentially expressed genes was analyzed using clusterProfiler^23^. For Gene Ontology (GO) Biological Process enrichment, gene sets were defined based on the direction of differential expression (positive or negative log_2_ fold-change) and the FDR significance threshold, with *Nfya* excluded from all input gene lists. Gene symbols were mapped to Entrez identifiers using clusterProfiler::bitr in conjunction with org.Mm.eg.db. GO enrichment analysis was performed using enrichGO (ont = “BP”, Benjamini–Hochberg correction). Enrichment results were optionally filtered to remove overly broad terms (gene count > 100). Redundant GO terms were reduced using semantic similarity–based simplification (clusterProfiler::simplify, Wang similarity). Selected GO terms were visualized using dot plots.

Reactome pathway enrichment was performed using ReactomePA::enrichPathway with mouse-specific annotations, applying analogous filtering and visualization strategies.

### Tissue processing

Mice were euthanized by ketamine/xylazine overdose, followed by blood collection, perfusion with ice-cold PBS containing 5 mM EDTA, and organ harvesting for flow cytometry or immunostaining. Cell suspensions were washed, centrifuged, and resuspended in Hanks’ balanced salt solution (HBSS Mg^2+^/Ca2^+^-free) supplemented with 0.06% BSA and 0.3 mM EDTA. BM was isolated by flushing femurs with 5 ml of HBSS using a 21-gauge needle. Aortic arches and hearts were embedded unfixed in Tissue-Tek O.C.T. compound. Mouse atherosclerotic plaques were enzymatically digested in 1 ml RPMI medium containing 10% fetal calf serum and 1.25 mg/mL Liberase at 37 °C for 1 hour.

### Flow cytometry

Red blood cells were lysed by incubation in ammonium chloride buffer (150 mM NH_4_Cl, 10 mM KHCO_3_, 0.1 mM Na_2_EDTA) for 5 minutes at room temperature. Leukocytes were stained in staining buffer for 20 minutes at 4 °C with antibodies against CD45 (30-F11), CD11b (M1/70), Gr1 (RB6-8C5), CD115 (AFS98),and F4/80 (BM8) (Biolegend).

For apoptosis analysis, cells were stained with annexin V–FITC according to the manufacturer’s instructions (Thermofisher, 88-8005-72). For proliferation analysis, mice were injected intraperitoneally with 1 mg of 5-bromo-2′-deoxyuridine (BrdU) 24 hours before sacrifice. BrdU incorporation was assessed in isolated cells by flow cytometry using the BrdU Flow Cytometry Kit (BD Pharmigen, 552598).

Flow cytometry was performed with a BD FACS Canto instrument, and data were analyzed using FlowJo software (BD Pharmigen).

### Immunostaining

Mouse tissues were fixed in 4% paraformaldehyde in PBS, paraffin-embedded, and sectioned at 5 µm thickness. Sections were stained with hematoxylin and eosin for histological assessment. For femoral artery wire injury experiments, thrombus-free tissue sections were used for subsequent analyses. In atherosclerotic lesions, the necrotic core was defined as the area devoid of nuclei underneath a formed fibrous cap. Total collagen content was assessed by Pricrosirius Red staining in consecutive sections.

For immunofluorescence staining, antigens were retrieved with citrate buffer (10mM, pH 6.0), followed by blocking with 5% goat serum in PBS. Sections were incubated overnight at 4 °C with rabbit anti-mouse CD68 (Abcam; 1:200) and mouse anti-mouse smooth muscle actin (SMA)–FITC (Sigma; 1:500). After extensive washing, sections were incubated with appropriate secondary antibodies conjugated to DyLight 488, DyLight 550, or DyLight 650 (Thermo Fisher; 1:500) and counterstained with DAPI (Sigma, D9542) to visualize nuclei.

Immunofluorescence images were acquired using an Evident VS200 Slide Scanner (Olympus). Histological and immunofluorescence sections were quantified by an investigator blinded to experimental condition using computer-assisted morphometric analysis with ImageJ software (National Institutes of Health).

### Statistics

Statistical analyses were conducted with GraphPad Prism 10. Data normality was assessed using the D’Agostino-Pearson omnibus test. Comparisons between two groups were by two-tailed unpaired Student *t*-test. Differences were considered statistically significant at a p value < 0.05. Data are expressed as mean ± SEM.

### Data and code availability

The data supporting the findings of this study are available from the corresponding authors upon reasonable request.

### Declaration of generative AI and AI-assisted technologies

ChatGPT 5.2 was used during the generation and improvement of the R code. ChatGPT 5.2 and DeepL write were used to improve English grammar. After using these tools, the authors reviewed and edited the content as needed and take full responsibility for the content of the publication.

## Results

### NF-Y subunit expression is associated with distinct transcriptional states in myeloid cells

To investigate NF-Y transcriptional regulation in myeloid cells, we generated an integrated single-cell transcriptomic dataset combining atherosclerotic lesional cells from published studies in mouse models of hypercholesterolemia with our own analysis of BM, blood, and lesional cells from *Apoe*^*-/-*^ mice fed a HFD for 16 weeks (see Methods). Using this integrated dataset, we assessed the expression of the three main NF-Y subunits (*Nfya, Nfyb*, and *Nfyc*) across key myeloid cell populations—monocytes, neutrophils, and macrophages—in different anatomical compartments. In BM and blood neutrophils, all three subunits were expressed along the maturation trajectory from progenitor to mature cells, with higher expression observed at the progenitor stage (**Figure S1A**), consistent with previous evidence linking NF-Y activity to proliferative states^6^. Monocytes likewise expressed all these transcripts in both BM and blood, with no marked differences between *Ly6c2*^*high*^ and *Ly6c2*^*low*^ monocyte subpopulations (**Figure S1B-D**). Although the proportion of cells expressing NF-Y subunits was highest in the BM, transcripts were also detected in blood and atherosclerotic lesions, consistent with NF-Y expression in differentiated myeloid cells. In vascular lesions, NF-Y subunit expression was observed across myeloid populations and other lesional cell types, both stromal and hematopoietic, with T-cell subpopulations showing the highest expression (**Figure S1E**). Together, these analyses demonstrate that differentiated myeloid cells express NF-Y subunit transcripts, with expression varying with cell subpopulation and anatomical location, consistent with our previous protein-level observations^14^.

We next focused on lesional macrophages by subsetting this population from the integrated dataset. Dimensionality reduction and unbiased clustering identified seven macrophage clusters (**Figure 1A**), defined by the expression of established marker genes: *Spp1* (Spp1 macrophages, Cluster 2), *Il1b* (pro-inflammatory Il1b+ macrophages, Clusters 1 and 4), *Stmn1* and *Smc4* (cycling macrophages, Cluster 3) and *Folr2, Retnla*, and *Ltc4s* (resident-like macrophages, Clusters 4, 6, and 7) (**Figure 1B**). Analysis of *Nfya* expression across clusters revealed variable expression, without enrichment in a single macrophage cluster (**Figure 1C**). We therefore stratified macrophages based on relative *Nfya* expression and compared transcriptional profiles between *Nfya*^*high*^ and *Nfya*^*low*^ populations (see Methods). *Nfya*^*high*^ macrophages showed elevated expression of genes associated with cell proliferation, including *Mki67*, multiple histone genes (*Hist1h4d, Histo1h1e*, and *Histh2ap*), and the integrator complex subunit *Ints6* (**Figure 1D**). In contrast, *Nfya*^*low*^ macrophages were enriched for transcripts linked to lipid handling and phagocytic function, such as *Cd5l* and *Gpnmb*^24, 25^, as well as *Mgp*, involved in extracellular matrix interactions, and inflammatory mediators including *Ccl5* and *S100a8*. Consistent with these findings, transcriptional signature analysis indicated enhanced phagocytic and metabolic programs in *Nfya*^*low*^ macrophages (**Figure 1E**). Together, these data indicate that relative *Nfya* expression distinguishes proliferative versus phagocytic transcriptional states among lesional macrophages.

**Figure 1.**
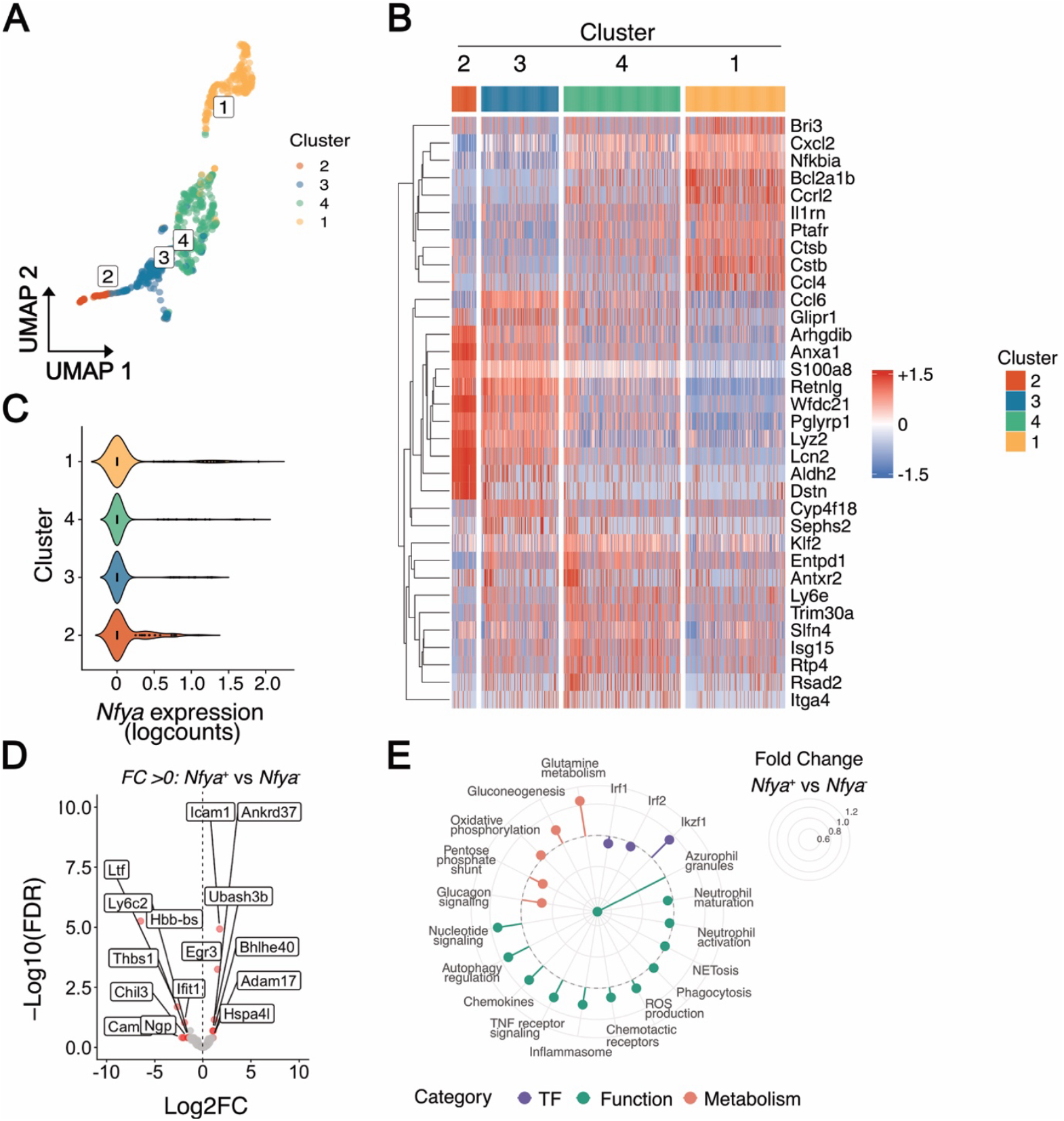
*Nfya* expression is associated with distinct transcriptional states in lesional macrophages. Single-cell transcriptomic analysis of macrophages identified in mouse atherosclerotic lesions in an integrated multi-study dataset (see Methods). **(A)** Uniform Manifold Approximation and Projection (UMAP) of lesional macrophages. **(B)** Heatmap of scaled, log-normalized gene expression across identified macrophage clusters. **(C)** Log-normalized *Nfya* expression across macrophage clusters. **(D)** Volcano plot showing differentially expressed genes between *Nfya*^*high*^ and *Nfya*^*low*^ macrophages. **(E)** Circulator lollipop plot showing selected differentially expressed genes distinguishing *Nfya*^*high*^ *and Nfya*^*low*^ macrophages.

We extended this analysis to lesional neutrophils and focused on this population by performing dimensionality reduction and unbiased clustering. We identified four clusters (**Figure 2A**), corresponding to immature neutrophils (Cluster 2), transitioning neutrophils (Cluster 3), interferon-activated neutrophils (Cluster 4), and activated mature neutrophils (Cluster 1) (**Figure 2B**). The proportion of *Nfya*-expressing cells was increased in Cluster 2 (**Figure 2C**). Interestingly, *Nfya* expression was also detected at the site of production within the BM (**Figure S2A, B**). A separated analysis of BM and blood identified 15 clusters, where BM neutrophil progenitors (Cluster 15) showed the highest proportion of *Nfya*-expressing cells, although total *Nfya* counts per cell increased as cells matured (**Figure S2B**). Compared with *Nfya*^*low*^ neutrophils, *Nfya*^*high*^ neutrophils displayed higher expression of transcripts associated with activation, such as *Adam17* (**Figure 2D**), along with an enhanced chemotaxis signature (**Figure S2C**). In contrast, *Nfya*^*low*^ neutrophils showed an immature phenotype, characterized by the expression of *Ltf, Camp* or *Ngp* (**Figure 2D**), as well as increased interferon signalling (**Figure 2D, Figure S2D**), which has previously been linked to a heightened activation state^26^. Similar to the differences observed in lesional neutrophils, circulating neutrophils also showed distinct transcriptional profiles between *Nfya*^*high*^ and *Nfya*^*low*^ neutrophils (**Figure S2E**). These results support the concept that *Nfya* expression levels are associated with distinct maturation and activation transcriptional profiles in neutrophils, both in circulation and upon recruitment to the atherosclerotic lesion.

**Figure 2.**
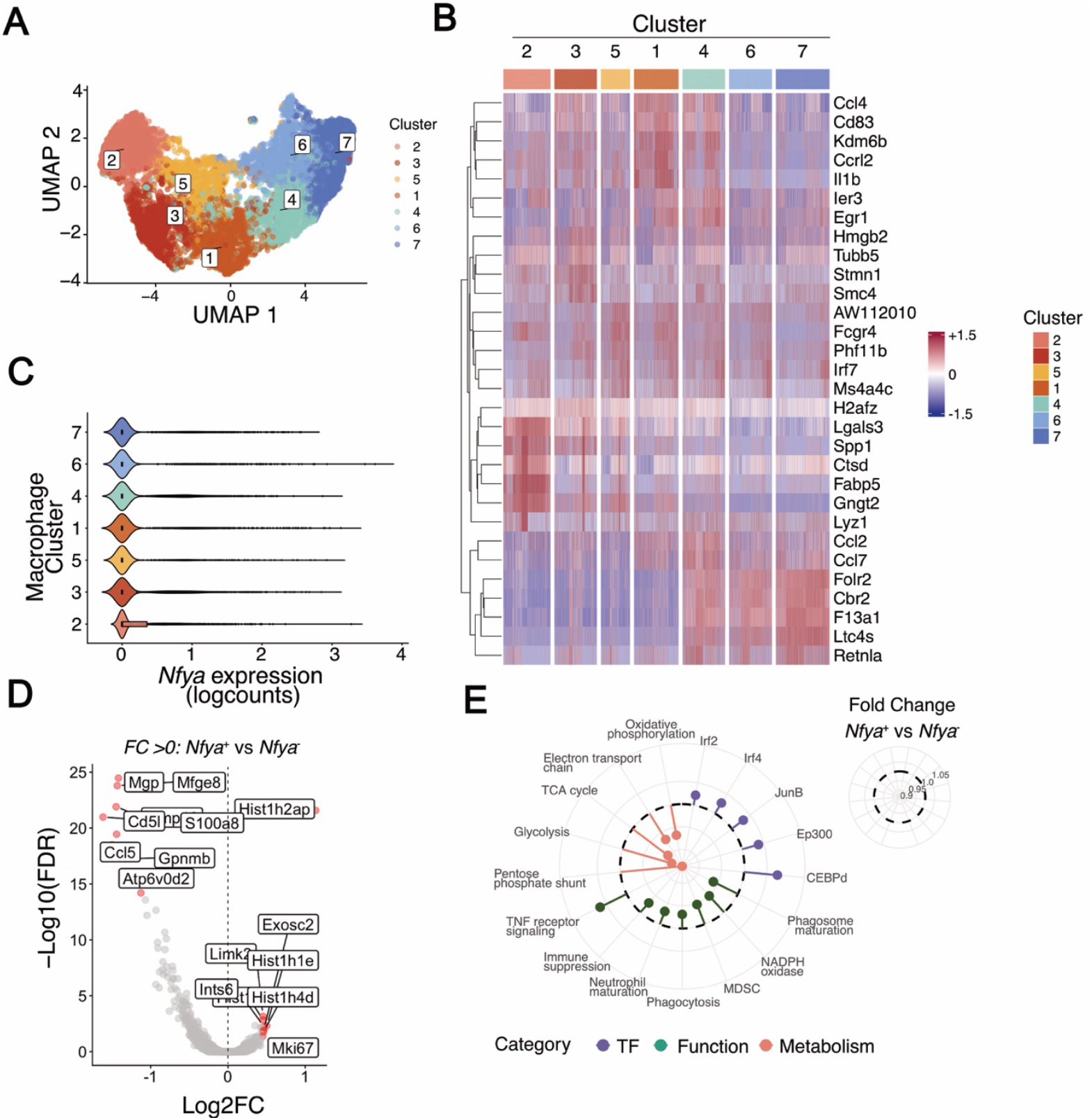
*Nfya* expression is associated with neutrophil maturation and activation states. Single-cell transcriptomic analysis of neutrophils identified in mouse atherosclerotic lesions in an integrated multi-study dataset (see Methods). **(A)** UMAP of lesional neutrophils. **(B)** Heatmap of scaled, log-normalized gene expression across identified neutrophil clusters. **(C)** Log-normalized *Nfya* expression across neutrophil clusters. **(D)** Volcano plot showing differentially expressed genes between *Nfya*^*high*^ and *Nfya*^*low*^ neutrophils. **(E)** Circulator lollipop plot showing enrichment of indicated transcriptional signatures in *Nfya*^*high*^ versus *Nfya*^*low*^ neutrophils.

### Myeloid-specific deletion of *Nfya* aggravates vascular disease

We next investigated whether myeloid-specific absence of NF-Y affects vascular disease progression *in vivo*. To do this, we generated mice with conditional deletion of the gene encoding the NF-Y A subunit (*Nfya*) by crossing *Nfya*^*fl/fl*^ mice with *LysM*^*CRE*^ transgenic mice and backcrossing this model onto the *Apoe*-deficient background. To ensure that *Nfya* deletion was confined to the hematopoietic compartment, BM cells were transplanted from *Apoe*^−*/*−^ *Nfya*^*fl/fl*^ or *Apoe*^−*/*−^ *LysM*^*CRE/+*^ *Nfya*^*fl/fl*^ donors into lethally irradiated *Apoe*^*-/-*^ recipients. Following hematopoietic reconstitution for 4 weeks, mice were fed a HFD for 12 weeks to induce atherosclerosis. Body weight did not differ between groups during the course of the diet (**Figure S3A**).

Flow cytometry analysis revealed reduced circulating neutrophil counts in mice with myeloid-specific *Nfya* deletion (**Figure S3B, F**), while total monocyte numbers were unchanged (**Figure S3C, G**). Within the monocyte compartment, the proportion of pro-inflammatory monocytes was reduced (**Figure S3D**), accompanied by an increase in non-classical monocytes (**Figure S3E**). Lymphoid populations were modestly increased in mice reconstituted with *Apoe*^−*/*−^ *LysM*^*CRE/+*^ *Nfya*^*fl/fl*^ BM (**Figure S3H-J**). In contrast, neutrophil and monocyte populations in the BM were comparable between groups (**Figure S3K, L**), indicating that the observed differences in circulating cells were not attributable to altered myelopoiesis.

Despite the reduction in circulating neutrophils and pro-inflammatory monocytes, mice with myeloid-specific *Nfya* deletion had an increased atherosclerotic burden. Lesion area was significantly increased in the aortic arch and thoracic aorta (**Figure 3A**), as well as in the aortic sinus, but not in the ascending aorta (**Figure 3B**). To determine whether these findings extended to other modes of vascular inflammation, we next used a femoral artery wire injury model of neointimal thickening in *Apoe*^−*/*−^ *LysM*^*CRE/+*^ *Nfya*^*fl/fl*^ mice fed a normal diet. Consistent with the atherosclerosis data, wire injury induced more pronounced neointimal thickening in *Apoe*^−*/*−^ *LysM*^*CRE/+*^ *Nfya*^*fl/fl*^ mice compared with *Nfya*^*fl/fl*^ controls, without significant differences in vessel stenosis (**Figure 3C**). Together, these results indicate that myeloid-specific loss of *Nfya* expression exacerbates vascular disease progression in response to both lipid-driven and injury-induced vascular injury.

**Figure 3.**
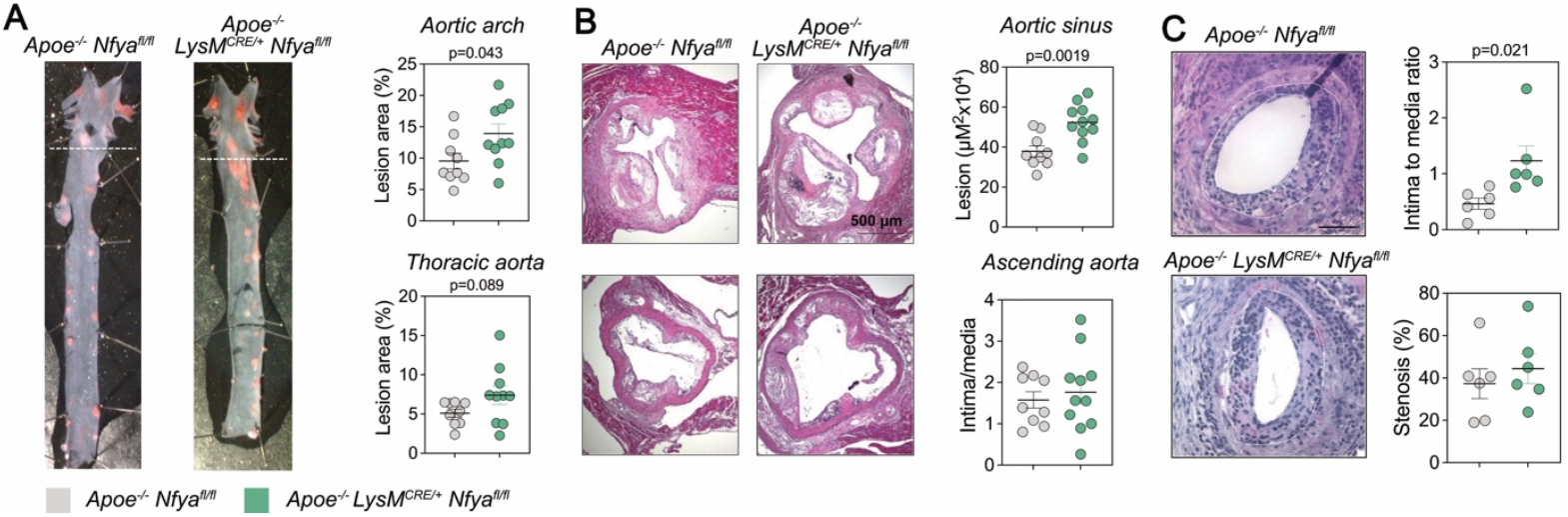
Myeloid-specific deletion of NF-YA increases atherosclerotic burden and injury-induced neointimal thickening. Lethally irradiated *Apoe*^−*/*−^ mice were reconstituted with bone marrow cells from either *Apoe*^−*/*−^ *Nfya*^*fl/fl*^ donors (control) or *Apoe*^−*/*−^ *LysM*^*CRE/+*^ *Nfya*^*fl/fl*^ donors (myeloid-specific *Nfya* ablation). Mice were fed a high-fat diet for 12 weeks (A, B) or normal diet (C). **(A)** Aortas were stained *en face* with Oil Red O to visualize atherosclerotic lesions (red). Representative images are shown. Graphs show quantification of lesional area in the aortic arch (top) and thoracic aorta (bottom), expressed as percentage of total area. **(B)** Cross-sections of the aortic sinus and ascending aorta were stained with hematoxylin and eosin. Representative images are shown (scale bar = 500 µm). Graphs show quantification of lesional area in the aortic sinus (top), and intima-to-media ratio in the ascending aorta (bottom). **(C)** Left and right femoral arteries of *Apoe*^−*/*−^ *Nfya*^*fl/fl*^ mice and *Apoe*^−*/*−^ *LysM*^*CRE/+*^ *Nfya*^*fl/fl*^ mice were subjected to wire injury. Representative images are shown (scale bar = 200 µm). Graphs show neointimal thickening (intima-to-media ratio, top) and vessel stenosis (bottom) quantified 7 days post-injury. Data are presented as mean ± SEM. Statistical significance was assessed by unpaired two-sided Student t-test.

### Myeloid *Nfya* deletion promotes features of plaque instability

To further assess the impact of myeloid-specific *Nfya* deletion on plaque composition, we analyzed advanced atherosclerotic lesions from the same cohort of BM-reconstituted mice. Quantification of collagen content revealed no significant differences between genotypes (**Figure 4A**). In contrast, necrotic core area was significantly larger in mice lacking *Nfya* in myeloid cells (**Figure 4B**), consistent with enhanced lesional cell death. Immunohistochemical analysis showed a reduction in SMA^+^ smooth muscle cells (SMC) within atherosclerotic lesions, while the proportion of CD68^+^ macrophages was unchanged (**Figure 4C**). These findings indicate that myeloid-specific *Nfya* deletion is associated with features of plaque instability, including increased necrotic core formation and reduced SMC content, without overt changes in macrophage abundance.

**Figure 4.**
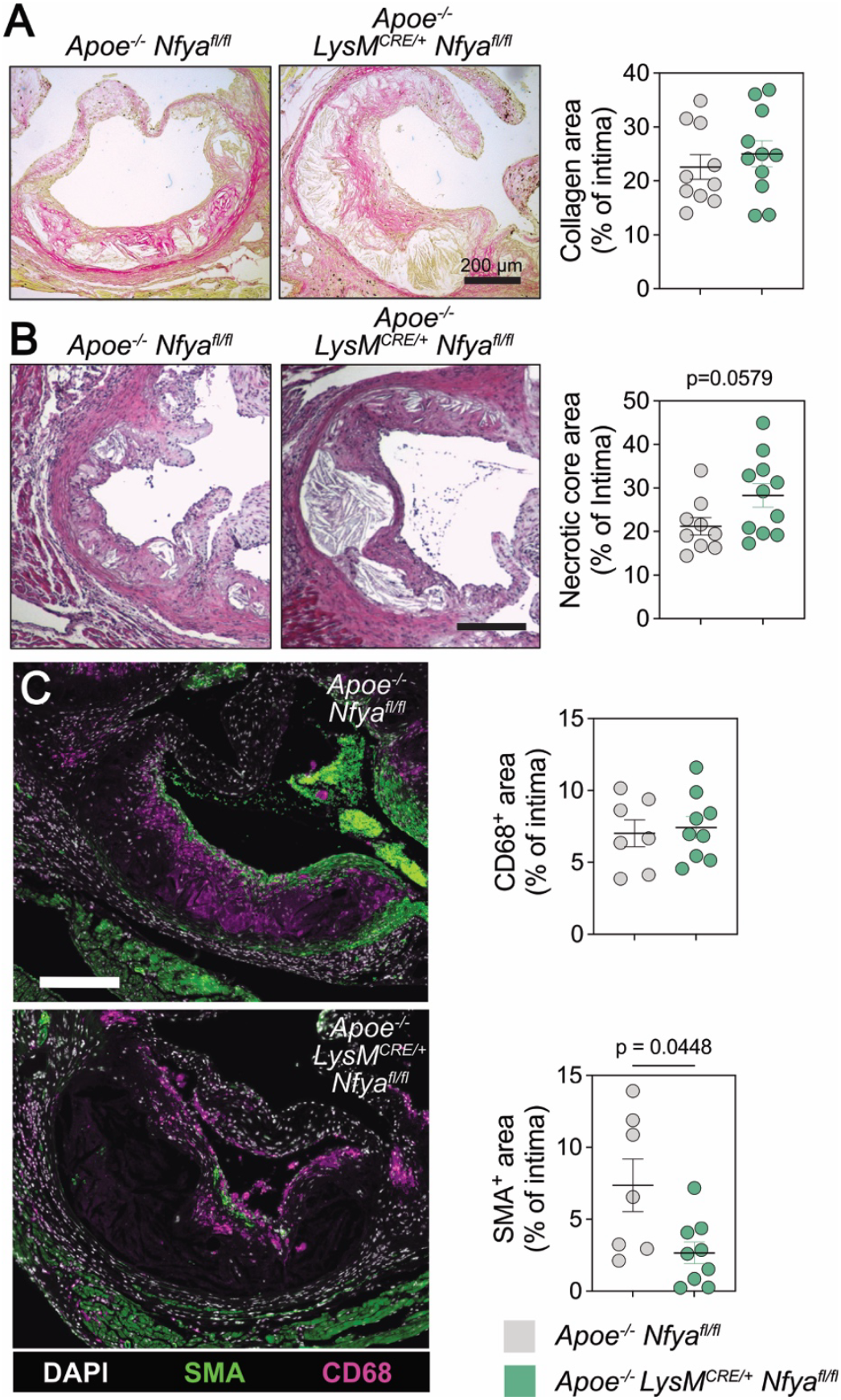
Myeloid-specific *Nfya* deletion is associated with features of plaque instability. Lethally irradiated *Apoe*^−*/*−^ mice were reconstituted with bone marrow from either *Apoe*^−*/*−^ *Nfya*^*fl/fl*^ donors (control; n = 9) or *Apoe*^−*/*−^ *LysM*^*CRE/+*^ *Nfya*^*fl/fl*^ donors (myeloid-specific *Nfya* deletion; n = 10) and fed a high-fat diet for 12 weeks. Atherosclerotic lesions analyzed in this figure were obtained from the same cohort of bone marrow–reconstituted mice shown in Figure 3A–B. **(A)** Representative images of Picrosirius red–stained aortic sections (scale bar = 200 μm). The graph shows quantification of collagen content expressed as a percentage of intimal area. **(B)** Representative images of hematoxylin and eosin–stained aortic sections (scale bar = 200 µm). The graph shows quantification of necrotic core area as a percentage of total lesion area. **(C)** Representative images of aortic sections showing immunostaining for SMCs (SMA; green) and macrophages (CD68; magenta), and counterstaining of nuclei (DAPI; white) (scale bar = 200 µm). Graphs show quantification of percentage of area positive for CD68 (top) and SMA (bottom) relative to total intimal area. Data are presented as mean ± SEM. Statistical significance was assessed by unpaired two-sided Student t-test.

### NF-YA is required for myeloid cell survival under inflammatory stress

Given the increased necrotic core size observed in lesions from mice with myeloid-specific *Nfya* deletion, we next asked whether NF-YA regulates myeloid cell recruitment, survival, or death under acute inflammatory conditions. To address this question, we used a model of acute thioglycolate-induced peritonitis. *LysM*^*CRE/+*^ *Nfya*^*fl/fl*^ mice and *LysM*^*+/+*^ *Nfya*^*fl/fl*^ littermate controls were injected intraperitoneally with 3% thioglycolate, and peritoneal exudates were analyzed 16 hours later. Total immune cell numbers in the peritoneal cavity were reduced in mice lacking myeloid Nfya (**Figure 5A**), driven primarily by a decrease in neutrophils (**Figure 5B**), whereas monocyte and macrophage numbers were not significantly altered (**Figure 5C, D**).

**Figure 5.**
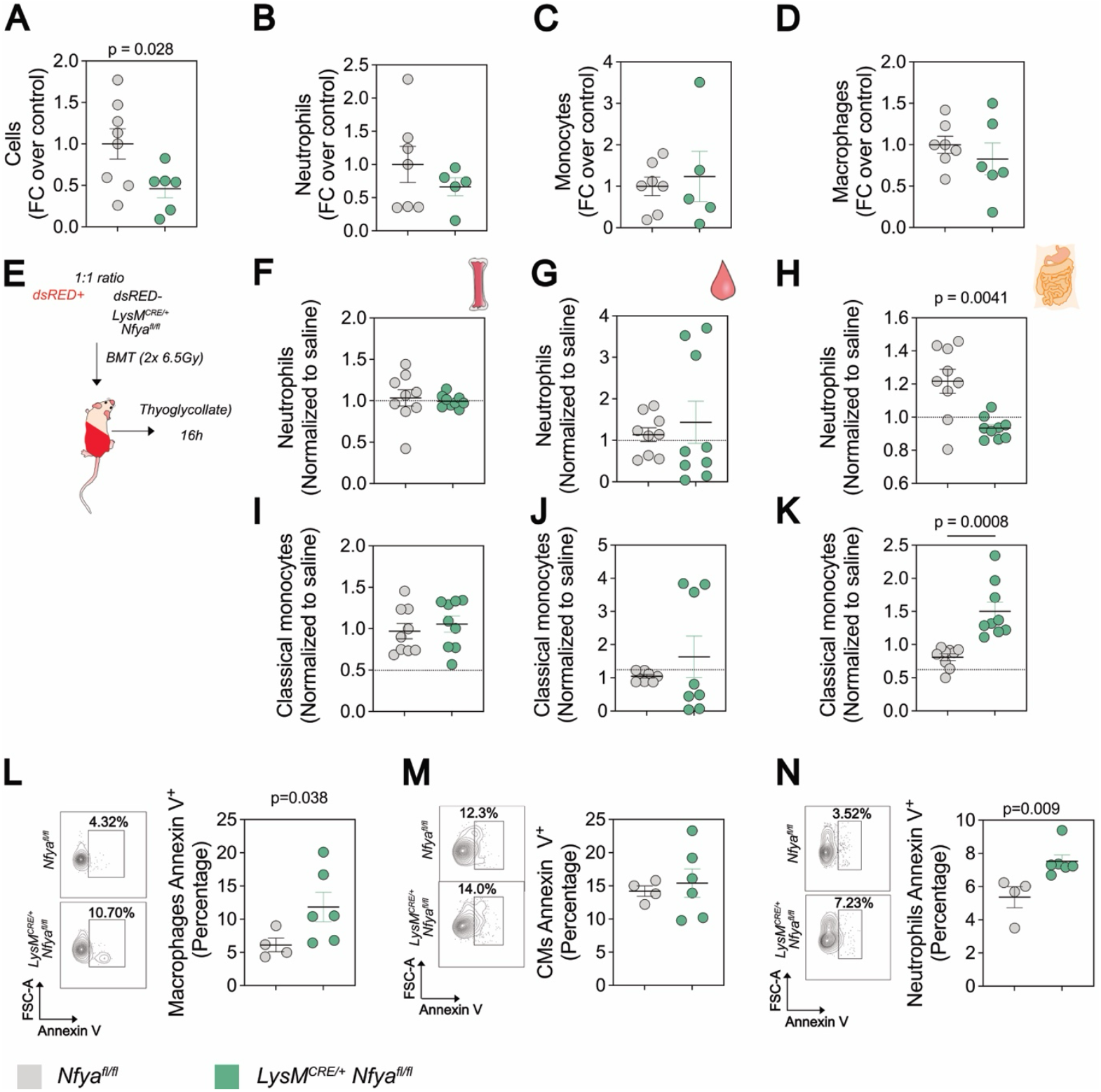
NF-YA is required for neutrophil and macrophage survival under inflammatory conditions. *LysM*^*+/+*^ *Nfya*^*fl/fl*^ (control; n = 7–8) and *LysM*^*CRE/+*^ *Nfya*^*fl/fl*^ (myeloid-specific *Nfya* deletion; n = 5–6) mice were injected intraperitoneally with saline or 3% thioglycolate to induce acute peritonitis. Myeloid cell populations were assessed 16 hours post-injection. **(A-D)** Relative counts of all cells **(A)**, neutrophils **(B)**, monocytes **(C)**, and macrophages **(D)** per mL of peritoneal exudate. **(E-K)** Female C57BL6/J mice were lethally irradiated and reconstituted with a 1:1 mixture of bone marrow cells from *Mafia*-GFP mice and *LysM*^*CRE/+*^ *Nfya*^*fl/fl*^ mice (n = 9 mice per group). After hematopoietic reconstitution, mice were injected intraperitoneally with 3% thioglycolate, and myeloid cell populations were assessed 16 hours later. (**F–H**) Flow cytometry analysis of neutrophils in bone marrow, blood, and peritoneal cavity, normalized to respective saline-injected controls. (**I–K**) Classical monocyte counts in bone marrow, blood, and peritoneal cavity. **(L-N)** Flow cytometry analysis of apoptotic cells (annexin V staining) in macrophages **(L)**, classical monocytes (CM) (**M**), and neutrophils **(N)** in peritoneal exudates. Representative plots (left) and quantification (right). n=4-6 mice/group. Data are presented as mean ± SEM. Statistical significance was assessed by unpaired two-sided Student t-test.

To determine whether these effects were cell-intrinsic, we generated competitive BM chimeras by reconstituting lethally irradiated recipient mice with a 1:1 mixture of dsRED^+^ *Nfya*-competent BM cells and *LysM*^*CRE/+*^ *Nfya*^*fl/fl*^ BM cells (*Nfya*-deficient, non-fluorescent) (**Figure 5E**). Following reconstitution and thioglycolate challenge, no significant between-genotype differences were found in neutrophil or monocyte numbers in the BM or blood (**Figure 5F, G, I, J**). In contrast, within the peritoneal cavity, neutrophils derived from *Nfya*-deficient cells were significantly less abundant (**Figure 5H**), whereas monocytes of *Nfya*-deficient origin were relatively increased (**Figure 5K**).

Analysis of cell death revealed increased apoptosis among *Nfya*-deficient neutrophils and resident macrophages, as assessed by annexin V staining, whereas classical monocytes were unaffected (**Figure 5L–N**). Notably, myeloid-specific *Nfya* deletion did not alter neutrophil or monocyte numbers in the BM or circulation under acute inflammatory conditions (**Figure S4A-D**), and nor did it affect monocyte or macrophage proliferation (**Figure S4E, F**) or macrophage uptake of acetylated low-density lipoprotein (**Figure S4G, H**). Together, these data indicate that NF-Y activity is required for the survival of neutrophils and macrophages under inflammatory stress, providing a mechanistic link between *Nfya* deletion, increased lesional cell death, and accelerated progression of atherosclerotic plaques.

## Discussion

In this study, we identify NF-Y as a transcriptional regulator that couples proliferative and survival programs to myeloid cell behavior during vascular inflammation. By integrating single-cell transcriptomics with myeloid-specific deletion of *Nfya* in models of aortic atherosclerosis, femoral artery endoluminal injury, and acute peritonitis, we provide evidence that NF-Y activity stratifies myeloid transcriptional states and preserves macrophage and neutrophil viability under inflammatory stress.

Our integrated single-cell analysis reveals detectable expression of NF-Y subunit transcripts across myeloid lineages in BM, blood, and atherosclerotic lesions. Although expression was more prominent in immature neutrophils and monocytes, NF-Y subunit transcripts were also detected in more mature cells in the circulation and inflamed tissue. Given the post-mitotic state of mature neutrophils, these findings suggest that NF-Y may regulate processes beyond cell-cycle control, including maturation-associated functions, chemotaxis, and cell activation. Consistent with this idea, *Nfya*-expressing neutrophils displayed differential expression of genes encoding granule proteins (*Ltf* (Lactoferrin), *Ngp* (Neutrophilic granule protein), and *Camp* (Cathelicidin)), adhesion molecules (*Icam1*), and interferon-responsive genes (*Ifit1*). In line with a role in cell migration or retention, our competitive peritonitis model showed reduced accumulation of *Nfya*-deficient neutrophils at the site of inflammation.

In lesional macrophages, cells with high *Nfya* expression were enriched for cell-cycle and biosynthetic signatures, including *Mki67* and histone genes, whereas macrophages with low *Nfya* expression displayed lipid-handling and phagocytic transcriptional features associated with foam cell formation. Notably, NF-YA deficiency in macrophages did not impair acetylated LDL uptake, indicating that NF-Y is unlikely to directly control lipid internalization. Instead, NF-Y may regulate downstream aspects of lipid handling or cellular stress adaptation, potentially limiting lipid toxicity in the inflammatory vascular environment.

A central functional finding of this study is that loss of NF-YA in myeloid cells increases apoptosis of neutrophils and resident macrophages during acute peritonitis and is associated with enlarged necrotic cores in atherosclerotic lesions. This survival phenotype is consistent with prior studies demonstrating that NF-Y is required for the survival of cycling hematopoietic stem cells (HSCs), but is dispensable in quiescent HSCs^6^, and that NF-Y deficiency impairs expansion of myeloid progenitors^8^. NF-Y has been shown to operate at the interface of proliferation and apoptosis by regulating genes involved in stress responses, including C/EBP homologous protein (CHOP), X-box binding protein 1 (XBP1), HIF-1α, and p53.^27, 28^ CHOP and XBP1, in particular, are key mediators of endoplasmic reticulum stress responses that enable cells to adapt to conditions such as misfolded protein accumulation, glucose deprivation, lipid toxicity, and hypoxia.^29^ Previous work has linked NF-Y deficiency to hepatic lipid accumulation and cell death, accompanied by altered expression of CHOP and XBP1.^30^

Our experimental design restricted *Nfya* deletion to *LysM*-expressing differentiated myeloid cells, thereby excluding direct effects in hematopoietic progenitors. Despite reduced circulating neutrophils and altered monocyte subset distribution in *Nfya*-deficient chimeras, these mice exhibited increased diet-induced atherosclerotic burden and enhanced neointimal thickening after mechanical injury. This apparent paradox underscores that disease progression is determined not only by circulating myeloid cell counts, but also by qualitative changes in cell survival, inflammatory responses, and tissue remodeling. Increased lesional cell death and necrosis may therefore be sufficient to amplify inflammation and drive disease progression.

Consistent with a broader protective role for NF-Y in vascular pathology, our previous work suggested that NF-Y also regulates SMC proliferation, potentially contributing to fibrous cap formation and plaque stability, while exacerbating luminal narrowing in restenosis. Distinguishing direct from indirect regulatory effects will require combining cell-type–specific perturbations with direct chromatin profiling of NF-Y targets in lesional macrophages, neutrophils, and SMCs. Integrating these data with established transcription factor networks governing myeloid maturation and effector functions may further clarify how NF-Y activity enables, constrains, or reshapes inflammatory gene programs in disease contexts.

Our data identify NF-Y as a transcriptional safeguard of macrophage and neutrophil survival during inflammation, with important consequences for plaque composition and vascular remodeling. By linking NF-Y activity to myeloid cell states and inflammatory pathology, this work expands current models of transcriptional control in myeloid biology and highlights NF-Y as a context-dependent regulator in cardiovascular disease. NF-Y dependence may be most critical under conditions of high biosynthetic demand, proliferative pressure, or inflammatory stress—hallmarks of myeloid cell recruitment and atherosclerotic plaque progression.

## Acknowledgments

We thank Sankar N Maity (Houston, Texas) for providing the Cbf^lfl/fl^ mouse and M.J. Andrés-Manzano for mouse genotyping.

## Sources of funding

Work in the CSR laboratory was supported by the Deutsche Forschungsgemeinschaft (CRC TRR332, projects A1 and A6; CRC1123, projects A6 and A7) and by the IZKF of the University of Münster. Work in the VA laboratory was supported by grant PID2022-141211OB-I00, funded by the Spanish Ministry of Science, Innovation, and Universities (MICIU) (MICIU/AEI/10.13039/501100011033) and by the European Regional Development Fund/European Union (EDRF/EU). The CNIC is supported by the Instituto de Salud Carlos III, the MICIU, and the Pro-CNIC Foundation and is a Severo Ochoa Center of Excellence (grant CEX2020-001041-S funded by MICIU/AEI/10.13039/501100011033). Work in the J.M.G.-G. laboratory was supported by grants PI24/00146, funded by the ISCIII and the Imas12.

## Disclosures

The authors declare no conflict of interest.

## SUPPLEMENTAL MATERIAL

**Figure S1.**
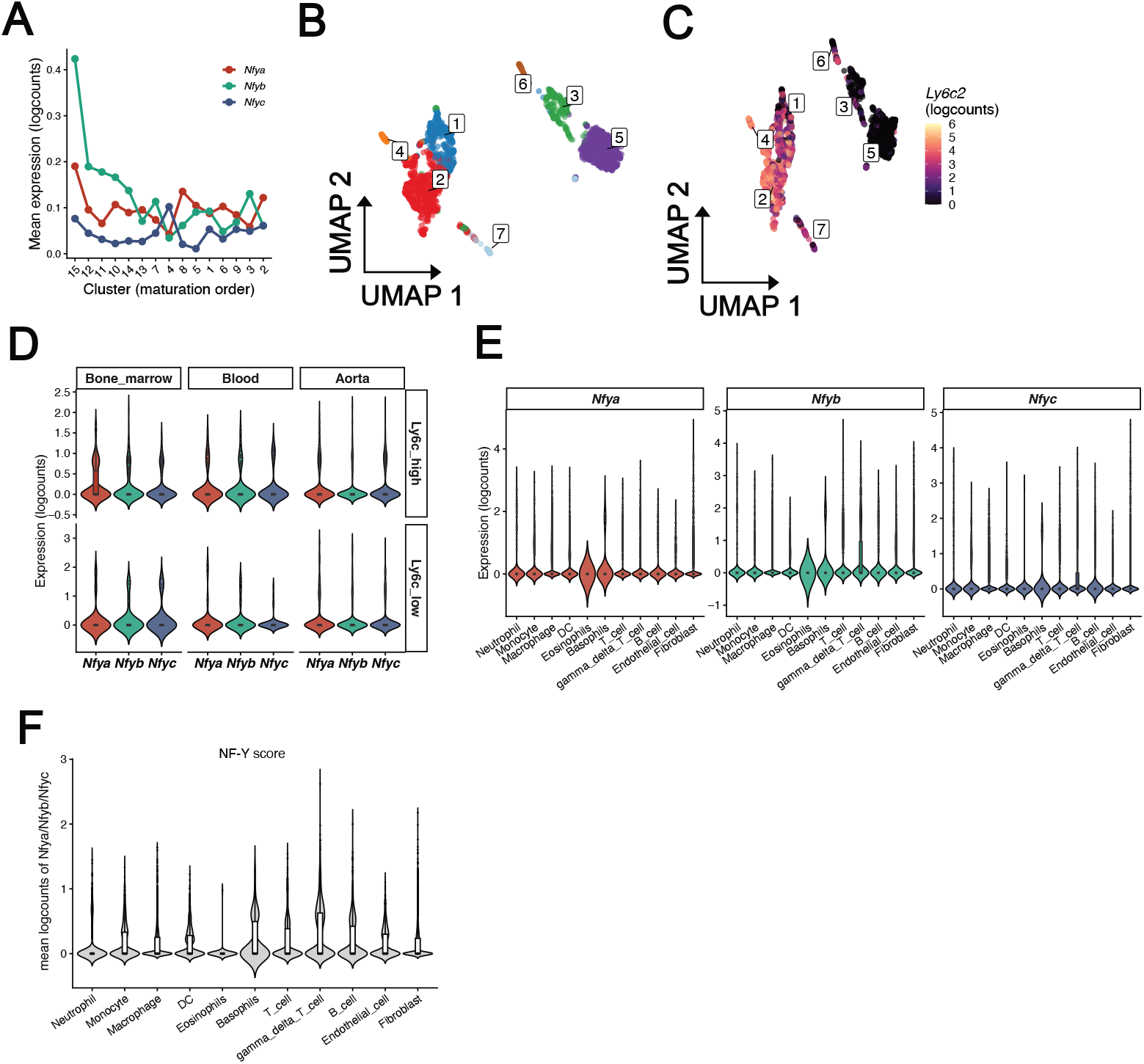
NF-Y subunit transcript expression across myeloid populations and anatomical compartments. **(A)** Log-normalized expression of *Nfya, Nfyb*, and *Nfyc* across indicated clusters of aggregated bone marrow and blood neutrophils from *ApoE*^*-/-*^ mice fed a high-fat diet for 16 weeks. **(B, C)** Uniform Manifold Approximation and Projection (UMAP) of monocytes colored by cluster identity **(B)** or by *Ly6c2* expression **(C). (D)** Expression of *Nfya, Nfyb*, and *Nfyc* in *Ly6c2*^*high*^ and *Ly6c2*^*low*^ monocyte subpopulations. **(E, F)** Expression of individual *Nfya, Nfyb*, and *Nfyc* subunits (E) or combined subunits (F) across hematopoietic and stromal cell populations within atherosclerotic lesions.

**Figure S2.**
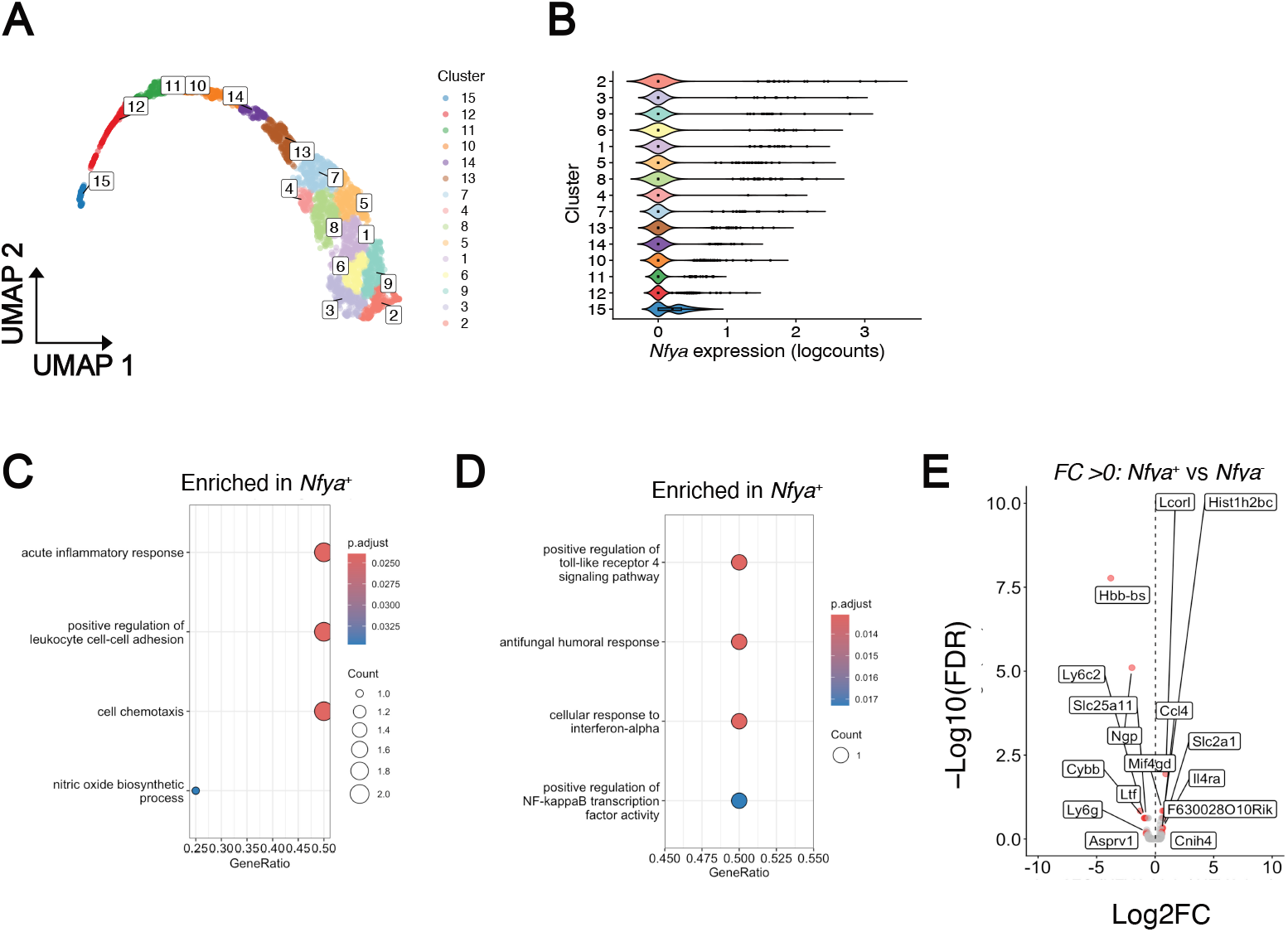
Transcriptomic features associated with *Nfya* expression in neutrophils across compartments. **(A)** Uniform Manifold Approximation and Projection (UMAP) of aggregated neutrophils from bone marrow and blood, colored by cluster. **(B)** Log-normalized *Nfya* expression across neutrophil clusters. **(C, D)** Gene ontology enrichment analysis of *Nfya*^*high*^ **(C)** and *Nfya*^*low*^ **(D)** lesional neutrophils from atherosclerotic lesions. **(E)** Volcano plot showing differentially expressed genes between *Nfya*^*high*^ and *Nfya*^*low*^ neutrophils in blood.

**Figure S3.**
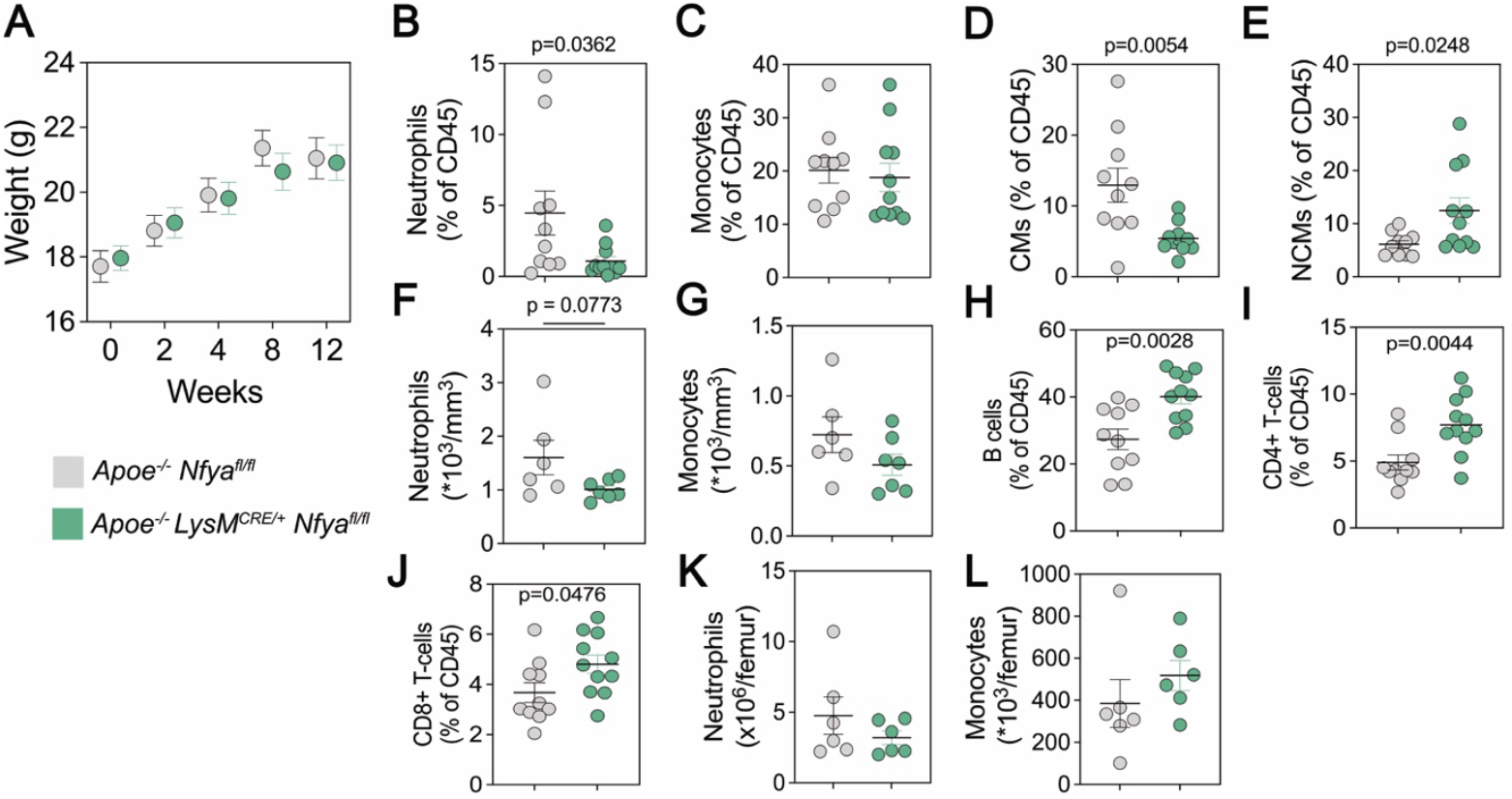
Effects of myeloid-specific Nfya deletion on body weight and circulating immune cell populations in hypercholesterolemic mice. Lethally irradiated *Apoe*^−*/*−^ mice were reconstituted with bone marrow from *Apoe*^−*/*−^ *Nfya*^*fl/fl*^ (n=9) or *Apoe*^−*/*−^ *LysM*^*CRE/+*^ *Nfya*^*fl/fl*^ (n=10) donors and fed a high-fat diet for 12 weeks. **(A)** Body weight measured at the indicated weeks following diet initiation. (**B–**J) Flow cytometry analysis of circulating immune cell populations: neutrophils (**B, F**; CD45^+^ CD11b^+^CD115^-^Gr-1^high^), total monocytes (**C, G**; CD45^+^ CD11b^+^CD115^+^), classical monocytes (**D**; CM, CD45^+^ CD11b^+^Gr1^high^ CD115^+^), non-classical monocytes (**E**; NCM, CD45^+^ CD11b^+^Gr1^high^ CD115^+^), B cells (**H**; CD45+ CD11b-B220+), CD4+ T-cells (**I**; CD45^+^ CD11b^-^CD4^+^), and CD8 T-cells (**J**; CD45^+^ CD11b^-^CD8^+^). **(K, L)** Flow cytometry analysis of bone marrow neutrophils **(K)** and total monocytes **(L)**. Data are presented as mean ± SEM. Statistical significance was assessed by unpaired two-sided Student t-test.

**Figure S4.**
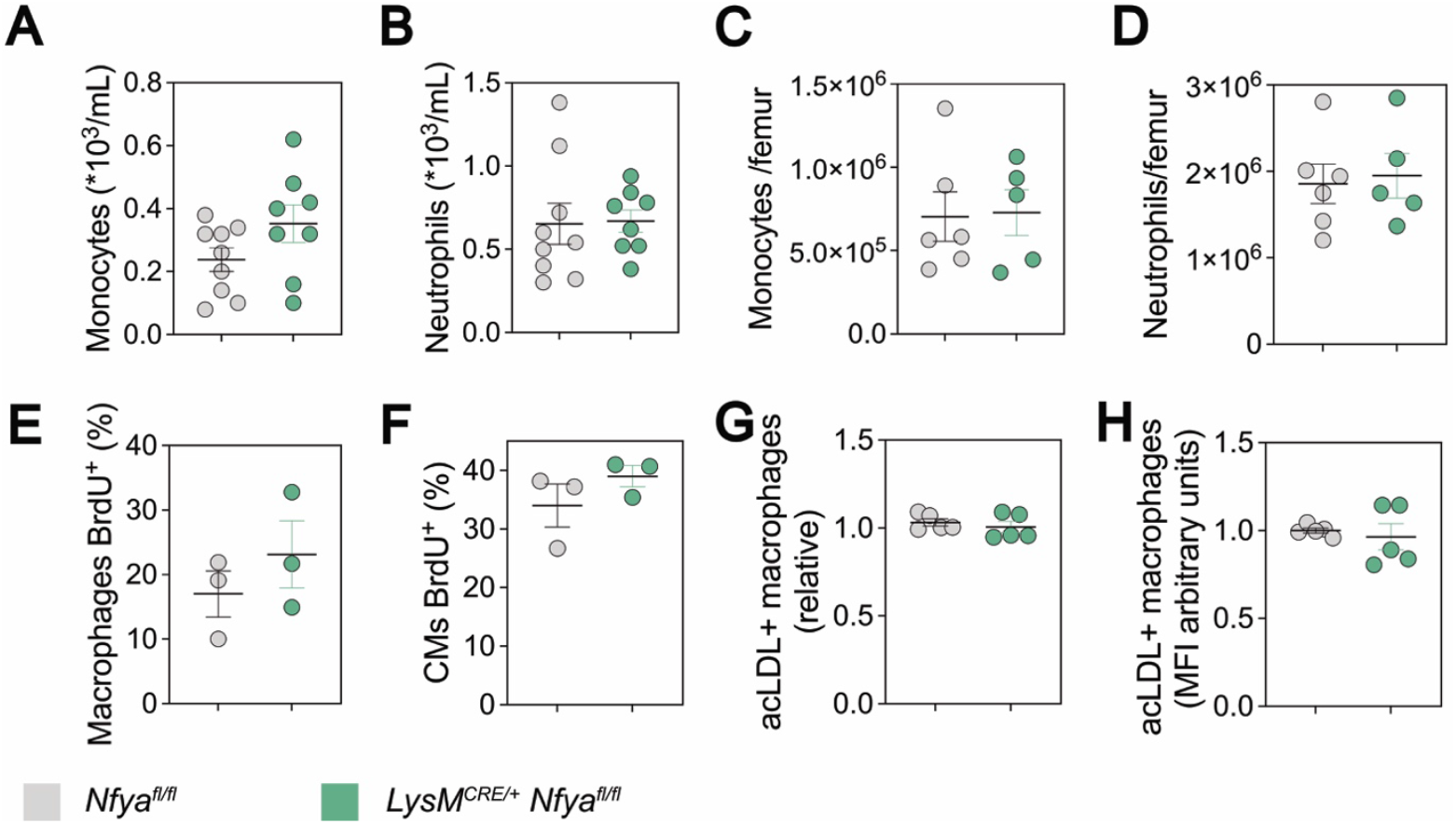
Myeloid-specific depletion of NF-YA does not alter bone marrow or circulating myeloid cells under acute inflammatory conditions. *Nfya*^*fl/fl*^ mice (control; n = 6–7) and *LysM*^*CRE/+*^ *Nfya*^*fl/fl*^ mice (n = 5–8) were injected intraperitoneally with saline or 3% thioglycolate to induce acute peritonitis, and myeloid cells were assessed after 16 hours. **(A, D)** Flow cytometry analysis of circulating monocytes **(A)**, circulating neutrophils **(B)**, bone marrow monocytes **(C)**, and bone marrow neutrophils **(D). (E, F)** Flow cytometry analysis of proliferating cells in peritoneal exudate, assessed by 5-bromo-deoxiuridine (BrdU) incorporation in macrophages **(E)** and classical monocytes (CM) (**F**). **(G, H)** Flow cytometry analysis of macrophage uptake of fluorescently labeled acetylated low-density lipoprotein (acLDL-FITC+), shown as percentage of acLDL^+^ cells **(G)** and mean fluorescence intensity (MFI) (**H**). Data are presented as mean±SEM. Statistical significance was assessed by unpaired two-sided Student t-test.

## Notes

### Competing Interest Statement

The authors have declared no competing interest.

